# Structural Basis of Complex Formation Between Mitochondrial Anion Channel VDAC1 and Hexokinase-II

**DOI:** 10.1101/2020.11.18.365965

**Authors:** Nandan Haloi, Po-Chao Wen, Qunlii Cheng, Meiying Yang, Gayathri Natarajan, Amadou KS Camara, Wai-Meng Kwok, Emad Tajkhorshid

## Abstract

Complex formation between hexokinase-II (HKII) and the mitochondrial channel VDAC1 plays a crucial role in regulating cell growth and survival; however, structural details of this complex remain elusive. We hypothesize that a conserved, hydrophobic helix (H-anchor) of HKII first inserts into the outer membrane of mitochondria (OMM) and then interacts with VDAC1 on the cytosolic leaflet of OMM to form a binary complex. To systematically investigate this process, we adopted a hybrid approach: 1) the membrane binding of HKII was first described with molecular dynamics (MD) simulations employing a membrane mimetic model with enhanced lipid diffusion, then 2) the resulting membrane-bound HKII was used to form complex with VDAC1 in millisecond-scale Brownian dynamics (BD) simulations. We show that H-anchor inserts its first 10 residues into the membrane, substantiating previous experimental findings. The insertion depth of the H-anchor was used to derive positional restraints in subsequent BD simulations to preserve the membrane-bound pose of HKII during the formation of the HKII/VDAC1 binary complex. Multiple BD-derived structural models were further refined with MD simulations, resulting in one stable complex. A major feature in the complex is the partial (not complete) blockade of VDAC1’s permeation pathway by HKII, a result supported by our comparative electrophysiological measurements of the channel in the presence and absence of HKII. Additionally, we showed how VDAC1 phosphorylation disrupts HKII binding, a feature that is verified by our electrophysiology recordings and have implications in mitochondria-mediated cell death.

## Introduction

Mitochondria are the primary source of ATP production in eukaryotic cells. Newly generated ATP is transported out of the mitochondrial matrix via the adenine nucleotide transporter at the inner mitochondrial membrane and out to the cytosol through the voltage-dependent anion channel (VDAC) located in the outer membrane of mitochondria (OMM). VDAC1, the most abundantly expressed isoform of VDAC, serves as the main conduit for the large flux of ions, ATP/ADP, metabolites, organic anions, and various respiratory substrates across the OMM^1–3^. In addition to its function as the major gateway in and out of mitochondria, VDAC1 also acts as a scaffold, regulating mitochondrial-triggered apoptotic signaling through interactions with a variety of proteins^2,4–6^.

VDAC1 has been reported to bind to pro- or anti-apoptotic proteins that can alter the permeability of the OMM and promote or prevent cell death^7–15^. For example, VDAC1 serves as a receptor for the cytosolic anti-apoptotic protein hexokinase (HK) which enhances cell survival^7,8^. Interaction with VDAC1 not only modulates cell survival but also gives HK preferential access to ATP for its catalytic activity of phosphorylating glucose to glucose-6-phosphate during glycolysis^16^.

HK is known as one of the primary factors of high glycolytic characteristics of rapidly growing tumor cell^17^. In particular, HK isoforms HKI and HKII are overexpressed in many types of cancers^4,18–20^. Elevated levels of HKI and HKII lead to a high rate of glycolysis, known as the Warburg effect, resulting in enhanced generation of lactic acid, a key component in promoting cell growth^21–23^. Coincidentally, HKI and HKII both differ from other HK isoforms with one extra structural element: in addition to two homologous hexose binding domains (N-domain and C-domain, Fig. 1A), HKI and HKII have a short N-terminal hydrophobic helix (termed H-anchor hereafter, Fig. 1A) that is thought to be capable of membrane-binding^24^. Given the importance of HK/VDAC1 interaction in cell growth and survival, disruption of this protein-protein complex has been identified as a potentially effective therapeutic strategy to prevent rapidly growing tumor cells^25–30^.

**Figure 1.**
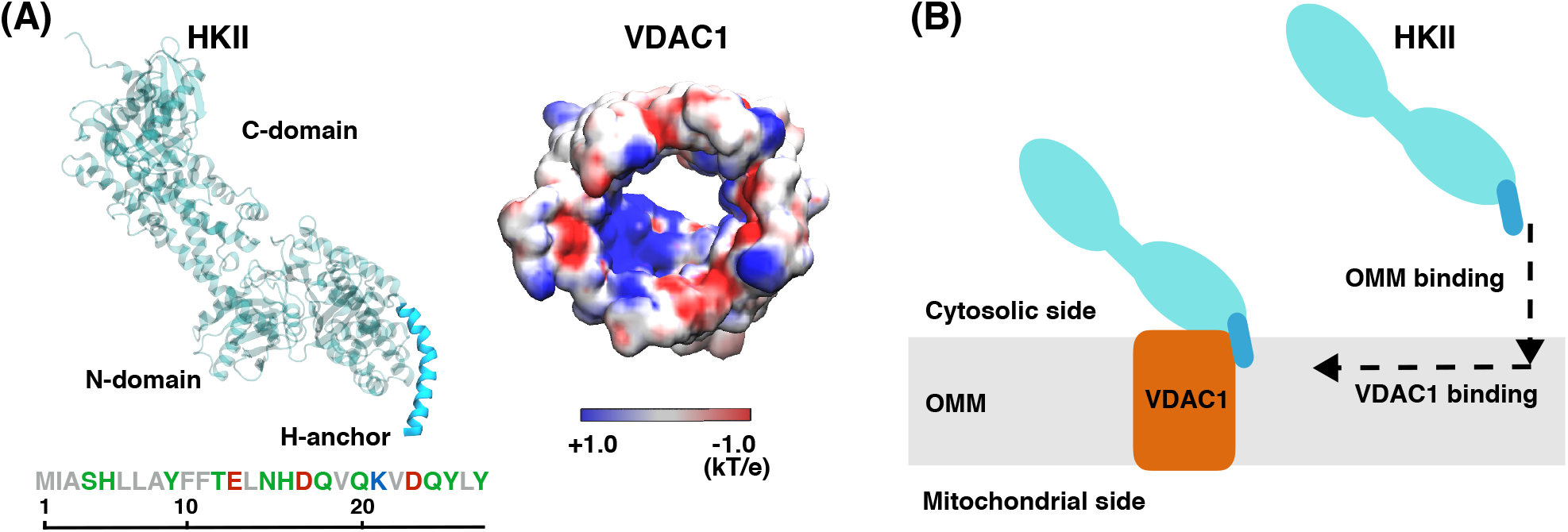
HKII and VDAC1 structures and the design of our modeling approach. (A, *Left*) The structure of HKII contains N- and C-domains (teal), and an N-terminal helix named here as H-anchor (cyan). The amino acid sequence of H-anchor is shown below, colored by their residue types: gray representing hydrophobic, green polar, red acidic, and blue basic amino acids. (A, *Right*) The top-down view of VDAC1 colored by its electrostatic potential, generated using the Poisson-Boltzmann (PB) equation solver in CHARMM-GUI^31–33^. (B) Schematic representation describing our modeling approach in which HKII binds the outer mitochondrial membrane (OMM) first, and then membrane-bound HKII forms a complex with VDAC1.

While a number of VDAC1 residues have been identified to be essential for interaction with HKI and HKII isoforms^7,21,34–37^, structural details of their complex remain unknown. Two protein-protein docking studies have been reported for modeling the complex formation between HK isoforms and VDAC1^38,39^. Both studies proposed direct plugging of the H-anchors of HKI/HKII into the pore of the barrel-shaped VDAC1. However, these models failed to address several key characteristics of HK/VDAC1 interactions, e.g., how the binding of HKI only partially reduces the maximal conductance of VDAC1^7,21,25,34^, as the plugged model would result in a complete blockade of the channel. Furthermore, at the docked interface, there is a mismatch between the hydrophobic H-anchor from HKI or HKII and the highly charged VDAC1 interior surface (Fig. 1A)^40^.

It is known that HKs can bind to membrane even in the absence of VDAC1^41–43^, and the H-anchor is shown to be essential for this membrane interaction^24,41,42^, likely due to its hydrophobic nature. Immunoblotting of mitochondria-bound HKI and HKII has shown that truncating the hydrophobic portions of the H-anchor disrupts HK binding to native OMM (even in the presence of VDAC1)^44,45^. The significance of the H-anchor in OMM binding is further emphasized by the fact that HK homologs lacking the H-anchor show no mitochondrial localization^20,46,47^. Possibly due to the inability to bind the membrane, truncation of the HKI H-anchor also eliminates the effect on the VDAC1 conductance^7^. Thus, the H-anchor in both HKI and HKII is vital for interaction with the OMM, a step which appears to be a prerequisite for HK-induced VDAC1 blockade. Henceforward, we hypothesized the following scenario in the complex formation between HKs (HKI or HKII) and VDAC1: first, HK binds to the OMM by inserting its H-anchor, a step that aligns the H-anchor for interaction with and binding to the outer-rim of VDAC1, in order to form the complex on the cytosolic surface of the OMM (Fig. 1B).

Based on the hypothesis described above, we designed a modeling approach involving two major steps. In the first step, we describe membrane binding of HKII with molecular dynamics (MD) simulations. Next, the resulting membrane-bound HKII is used to study the formation of its complex with VDAC1 using Brownian dynamics (BD) simulations. Our results show that HKII binds to the membrane by inserting its H-anchor, independent of VDAC1. The resulting complex reveals two key functional features of HKII/VDAC1 interaction: 1) a partial, and not complete, blockage of VDAC1 conductance through HKII interaction, and 2) how phosphorylation of VDAC1, and the corresponding phosphomimetic mutation, can disrupt the interaction between the two protein. Both of these predictions resulting from the computational model are supported by our designed electrophysiology measurements and mutagenesis experiments.

## Results and Discussion

### Spontaneous membrane binding and insertion of HKII

In order to investigate direct membrane interaction of HKII, we performed 10 independent 200-ns MD simulations using the highly mobile membrane mimetic (HMMM) model^48^ (Fig. 2A, more details in SI), during which binding of the N-domain of HKII (referred to as HKII-N) to lipid bilayers representing the OMM was simulated. Binding of HKII-N to the membrane was observed in 8 out of 10 replicas (Fig. 2B), all with the H-anchor inserting into the membrane. Spontaneous membrane binding occurs within the first 50 ns of the simulations in 7 out of the 8 successful replicates. After the first encounter with the membrane, the H-anchor is rapidly inserted into the membrane and the HKII-N remains membrane-bound for the remainder of the simulation (Fig. 2B). The H-anchor reaches a maximum depth of 10 Å below the bilayer’s phosphorus plane (as measured by the insertion of residue I2), penetrating well into the membrane’s hydrophobic core (Fig. 2B).

**Figure 2.**
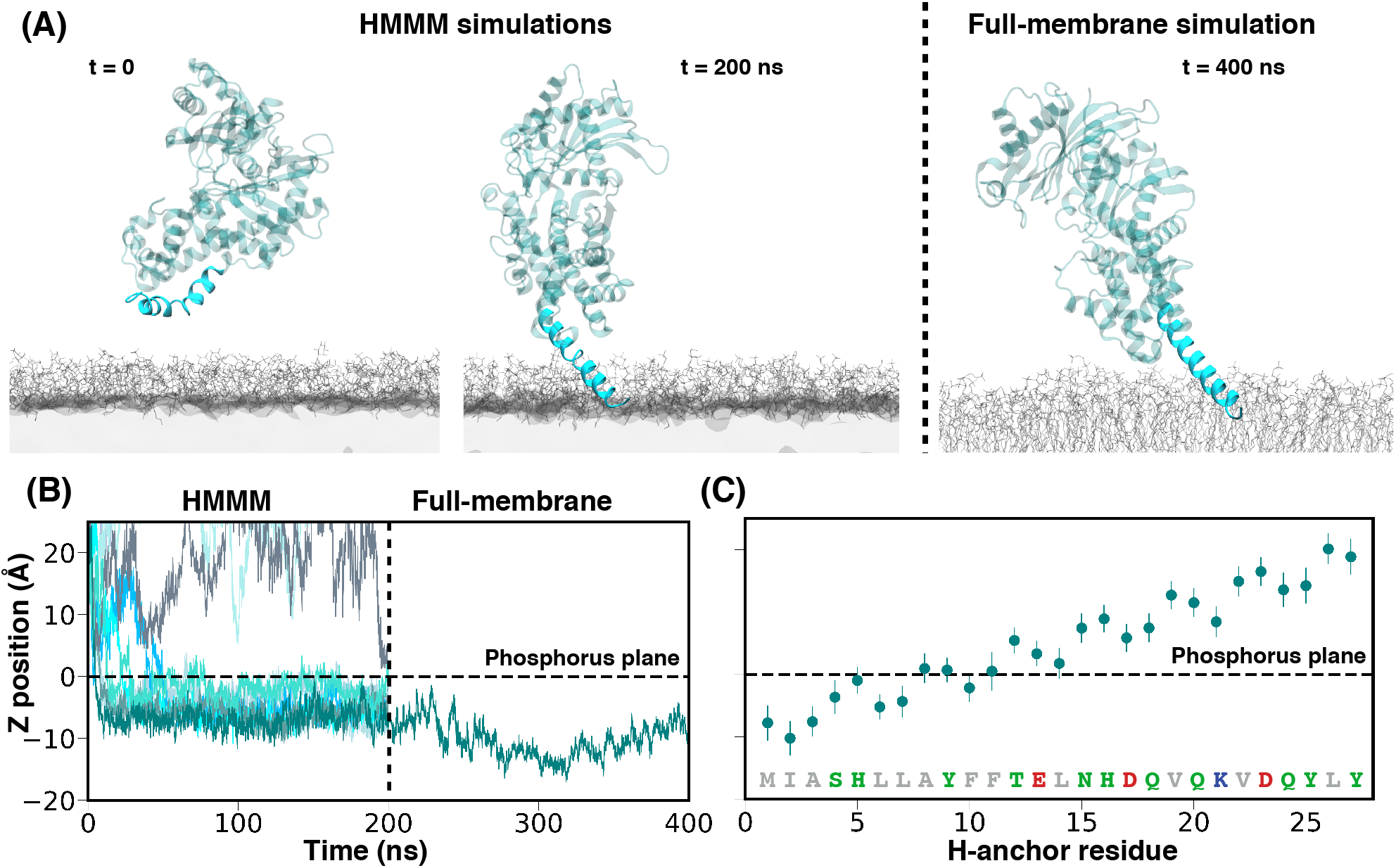
Membrane binding of HKII-N. (A) Initial and final, membrane-bound configurations of HKII-N in a representative trajectory. Initial, spontaneous membrane binding simulations (200 ns each) were performed using HMMM membranes (left panel). After membrane binding was achieved, the membrane was converted to a full membrane consisting of full-tailed lipids (right panel) and the system further simulated for 200 ns (totaling to 400 ns). The coloring scheme of HKII is identical to Fig. 1. Membranes are colored in gray. (B) Insertion depth of the protein into the membrane monitored by the position of I2 in HKII-N along the membrane normal (*z*-axis) with respect to the plane of the lipid phosphorus atoms (horizontal dashed line). The vertical dashed line separates the initial, HMMM membrane binding simulation (left side, all 10 simulation replicas shown), from the following 200-ns, full-membrane simulation performed for one of the systems (right side). (C) Mean and standard deviation of the relative *z* positions of the center of mass (C.O.M) of the H-anchor residues, calculated from the last 100 ns of the full-membrane simulation.

To ensure that the final membrane-bound model of HKII-N obtained from the HMMM simulations was stable, we converted one of the resulting membrane-bound systems to a conventional, full membrane and simulated it for additional 200 ns. The H-anchor remains buried into the hydrophobic core of the full membrane during the simulation (Fig. 2). In fact, the H-anchor appears to become even more engaged with the membrane during this additional simulation, as evidenced by deeper penetration of I2 into the membrane, reaching a maximum depth of 17 Å below the phosphorus plane of the lipid head groups’ approximately at *t* = 300 ns (Fig. 2B). These results provide strong support and a model for direct interaction of HKII with the membrane. Membrane partitioning of the H-anchor obtained from the full-membrane simulation shows that the first 10 residues remain stably bound and/or insert into the membrane (below the phosphorus plane) (Fig. 2C). This finding substantiates the experimental results where truncation of the first 10 residues from HKII was shown to disrupt its OMM binding^44^. Hence, our results provide the first structural model for membrane-bound HKII.

### HKII/VDAC1 complex formation

Once we established the membrane binding mode of HKII-N using MD simulations, we extended our study to investigate the molecular interaction between membrane-bound HKII-N with membrane-embedded VDAC1 by performing an aggregate of millisecond-scale atomic-resolution BD simulations. BD simulations were performed by placing 100 independent replicas of HKII-N around VDAC1 (Fig. 3A-C); each system was simulated for 20 *μ*s, totaling to 2 milliseconds. During the simulations, the membrane-bound pose of HKII-N was preserved using restraints designed based on the MD simulation results. These restraints allowed for the lateral diffusion of HKII-N around VDAC1 but prohibited significant vertical displacement of the system with respect to the membrane plane, thereby approximating the membrane-insertion depth and orientation of HKII-N (Fig. 3A). Details for the BD simulations are provided in SI text.

**Figure 3.**
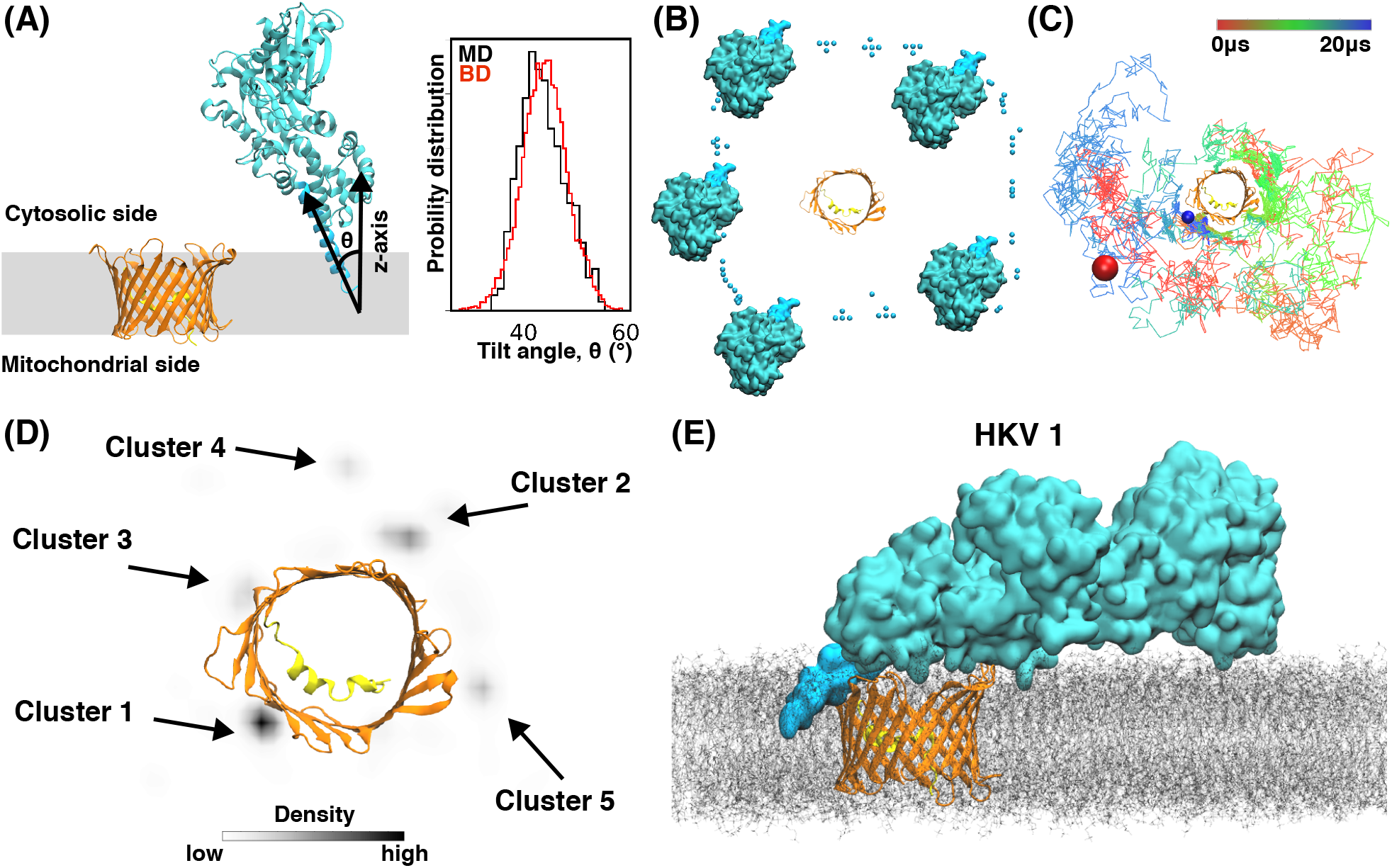
Binding of membrane-inserted HKII-N to VDAC1 and their complex formation. (A) Side view of the initial configuration in a BD simulation system. During the simulation, restraints along the *z*-axis (direction normal to the membrane plane represented as the gray box) were applied to the H-anchor, to maintain the positioning and the tilt angle distribution of the H-anchor (*Inset*) in BD simulations close to those observed in full-membrane MD simulations (see SI for more details). For tilt comparison, the distributions obtained from 1.5 *μ*s of ARBD simulations of HKII-N (in the absence of VDAC1) and the last 100 ns of full-membrane simulation of HKII-N are compared in the inset. VDAC1 is colored in orange with its N-terminal helix in yellow. The coloring scheme of HKII-N is identical to Fig. 1. (B) Top-down (cytosolic) view showing the initial placements of HKII-N around VDAC1 in independent BD simulations. Only 5 out of 100 HKII-N replicas are shown in full for clarity. The initial positioning of the other 95 replicas is represented by the C.O.M of the H-anchor. (C) A representative BD simulation trajectory showing the collision and interaction of HKII-N with different locations of the outer-rim of VDAC1 and its eventual binding. The colored line tracks the position of the C.O.M of the H-anchor over a 20-*μ*s BD simulation. The initial and final positions of the H-anchor are shown as red and blue spheres, respectively. (D) The density map of the C.O.M of H-anchor, calculated using the trajectories from all 100 independent BD replicates. The density map was calculated using the kernel density approximation module implemented in Seaborn^49^. Five distinct density hot-spots, each corresponding to a different cluster, are highlighted with black arrows. The clusters are ranked according to their population (Table S1). (E) A snapshot (at *t* = 350 ns) of membrane-embedded HKII/VDAC1 complex 1 (HKV1) during the full-membrane MD simulation.

The BD simulations revealed five distinct hot spots for HKII-N to bind VDAC1 while remaining anchored in the membrane (Fig. 3D). To further investigate whether these hot spots represent distinct HKII-N/VDAC1 complexes, we performed cluster analysis of HKII-N’s relative position to VDAC1 (using C_*α*_ root-mean-square deviation as a reference) for BD trajectories when the two are in close contact (*i.e.*, having at least 1 atom within 3.0 Å cutoff distance). Top five distinct clusters resulted from our analysis match the locations of the 5 BD hot spots (Fig. 3D). However, only Clusters 1-3 among the top 5 show direct contacts between the H-anchor and VDAC1 (Table S1). Since H-anchor is known to be crucial for HK/VDAC complex formation^7^, we considered only these three clusters for further refinement using MD simulations.

A representative HKII-N/VDAC1 complex model was selected from each of clusters 1–3 (termed HKV1, HKV2, and HKV3 hereafter) based on the strongest interaction energy between HKII-N and VDAC1 in the cluster. Each model was then used to generate a full-length HKII/VDAC1 complex by extending the C-domain from a terminal helix of HKII-N. The resulting complexes were then inserted into a fully-atomic membrane, creating three independent systems for additional MD equilibrations (Fig. 3E, S2). The contacts between the H-anchor and VDAC1 in HKV1 were maintained during the 350 ns refinement MD simulation (Fig. S2). Furthermore, a favorable interaction energy between H-anchor and VDAC1 during simulation indicated the stability of HKV1. The contacts between the H-anchor and VDAC1 in HKV2 were completely lost after 30 ns and did not re-form during the rest of the refinement MD simulation (Fig. S1). Though the H-anchor/VDAC1 contacts in HKV3 were maintained during the 350-ns MD simulation, these interactions were significantly weaker than those in HKV1 (Fig. S1). Given the importance of direct H-anchor/VDAC1 interactions in stabilizing the complex between HK and VDAC1^7^, we consider only HKV1 as the most relevant system for further structural analysis.

In HKV1, multiple hydrogen bonds and salt-bridge interactions between VDAC1 and HKII (H-anchor and N-domain) maintained the stability of the complex during the MD simulation (Fig. S3). Notably, no contacts were observed between the C-domain of HKII and VDAC1 (Fig. S3). This is consistent with previous experiments where mitochondrial binding of HKII (and HKI) was observed with immunoblotting of H-anchor+N-domain, but not with the C-domain only^45^.

### HKII modulates VDAC1 conductance

Upon complex formation, HKII covers a large fraction of the cytosolic surface of VDAC1 (Fig. 4A), primarily due to the contacts from the N-domain. This coverage results in an almost halved cytoplasmic opening of VDAC1 (reduced from 13.0 Å to 7.3 Å, Fig. 4B), suggesting that the VDAC1 conductance might be affected upon HKII binding. To further test this hypothesis and quantify the effect, we performed constant-voltage MD simulations for both HKII-free and HKII-bound states of VDAC1. Three independent simulations were performed for both states, each under a 50 mV membrane potential for 80 ns. Ionic currents obtained from these simulations showed that VDAC1 conductance is indeed partially reduced upon HKII binding (Fig. 4C,D), agreeing with previous electrophysiology measurements of HKI-bound VDAC showing partially reduced channel conductance^7,21,34^. On the other hand, the H-anchor plugged model proposed in previous computational studies^38,39^would virtually blocks the channel permeability completely, as indicated by the drastic narrowing of the pore to a 2.1 Å effective radius and a negligible current in constant voltage simulations (Fig. S4), which cannot account for the partial reduction of current observed experimentally upon HKII/VDAC binding.

**Figure 4.**
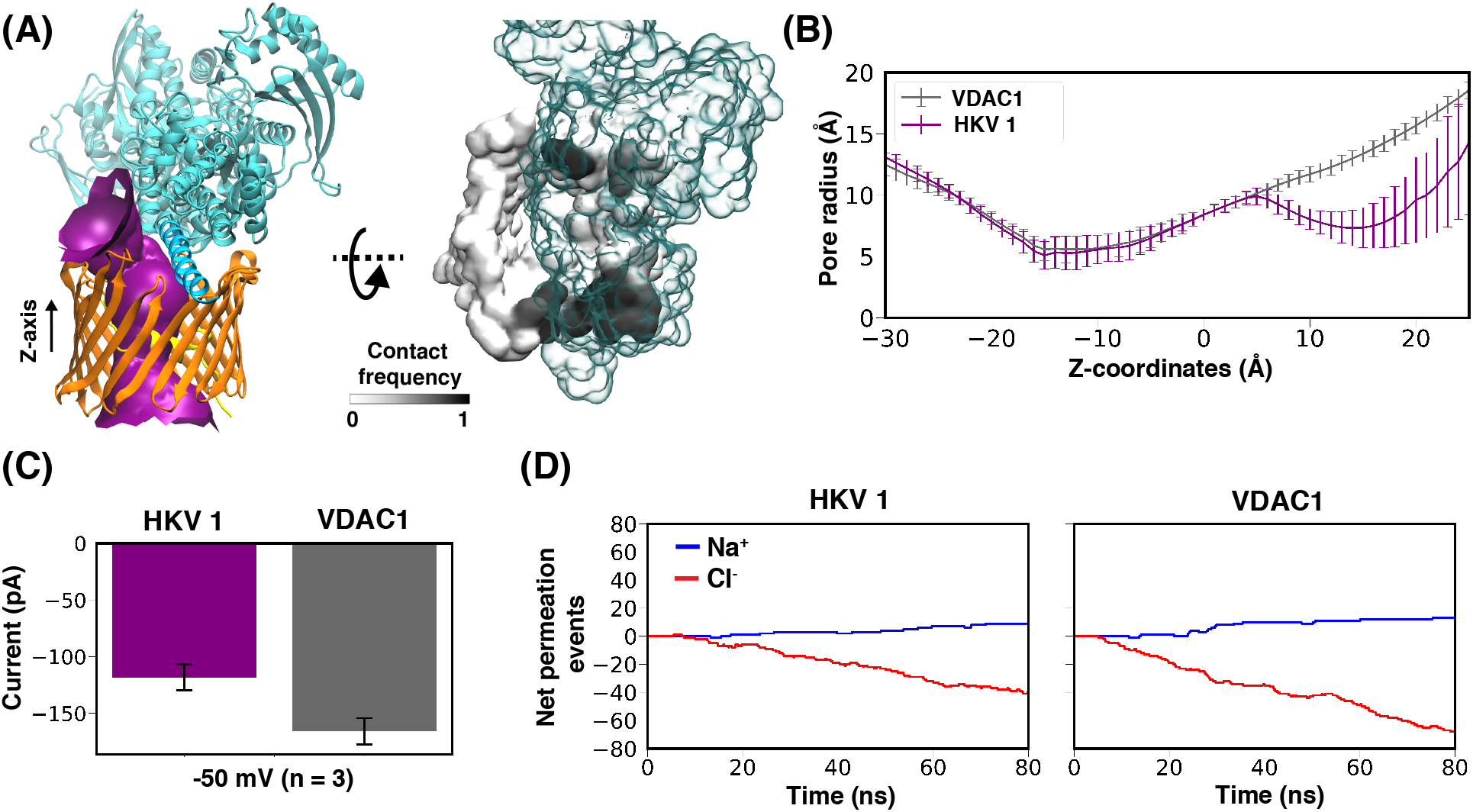
HKII/VDAC1 interaction in Complex HKV1. (A, *Left*) Purple surface showing VDAC1 pore radius profile in HKV1, made with the program HOLE^50^. Z-axis represents the membrane normal. (A, *Right*) Top-down (cytosolic) view of contact frequencies of HKII (transparent representation) mapped onto the surface of VDAC1 (black and white), calculated using the last 150 ns of MD simulation during which HKII/VDAC1 contacts converge (Fig. S2). (B) The average radius profile of the VDAC1 pore calculated using HOLE^50^ for both VDAC1 alone, and its complex with HKII (Complex HKV1). (C) The *in silico* VDAC1 current at −50 mV for a membrane-equilibrated VDAC1 alone or the VDAC1/HKII HKV1 complex calculated from electric-field MD simulations. The mean and standard deviation of current for each systems were calculated from 3 independent simulations. (D) The cumulative net number of channel-crossing events by Cl^−^ (red traces) and Na^+^ (blue traces), tracked over the time course of a representative MD simulation for VDAC1 or the HKV1 complex.

The hallmarked partial blockage of VDAC1 current upon HKII binding demonstrated in the simulations was further corroborated with a series of electrophysiology measurements. Using the planar lipid bilayer method previously described^51^, the reconstitution of wild type VDAC1 and its insertion into the bilayer were evidenced by its unique voltage-dependent gating characteristic in response to a voltage ramp protocol from 80 to +80 mV, where the maximal conductance occurs approximately between 50 and +50 mV and lower conductances at the more hyperpolarizing or depolarizing potentials (Fig. 5A). After washing away remaining soluble VDAC, current was monitored under a 30 mV test potential for 30 s at two minute intervals. When HKII was added to the *cis* chamber of the planar lipid bilayer (corresponding to the cytosolic side), there was a marked decrease in the maximal conductance of VDAC1 (Fig. 5B, S5). Steady-state levels were reached in approximately 6 minutes, and this latency may be due to the time it takes for the membrane insertion of HKII and its diffusion at the bilayer before reaching VDAC1. Upon reaching steady state, the effect of HKII on VDAC1 was stable for the subsequent 18 minutes of experiment time. Note that the addition of HKII resulted in only the reduction of VDAC1 conductance but not a complete blockade. These results strongly support the HKV1 model in MD simulations (Fig. 4) where the N-domain of HKII only partially covers the permeation pathway of VDAC1 and partially reduces the conductance.

**Figure 5.**
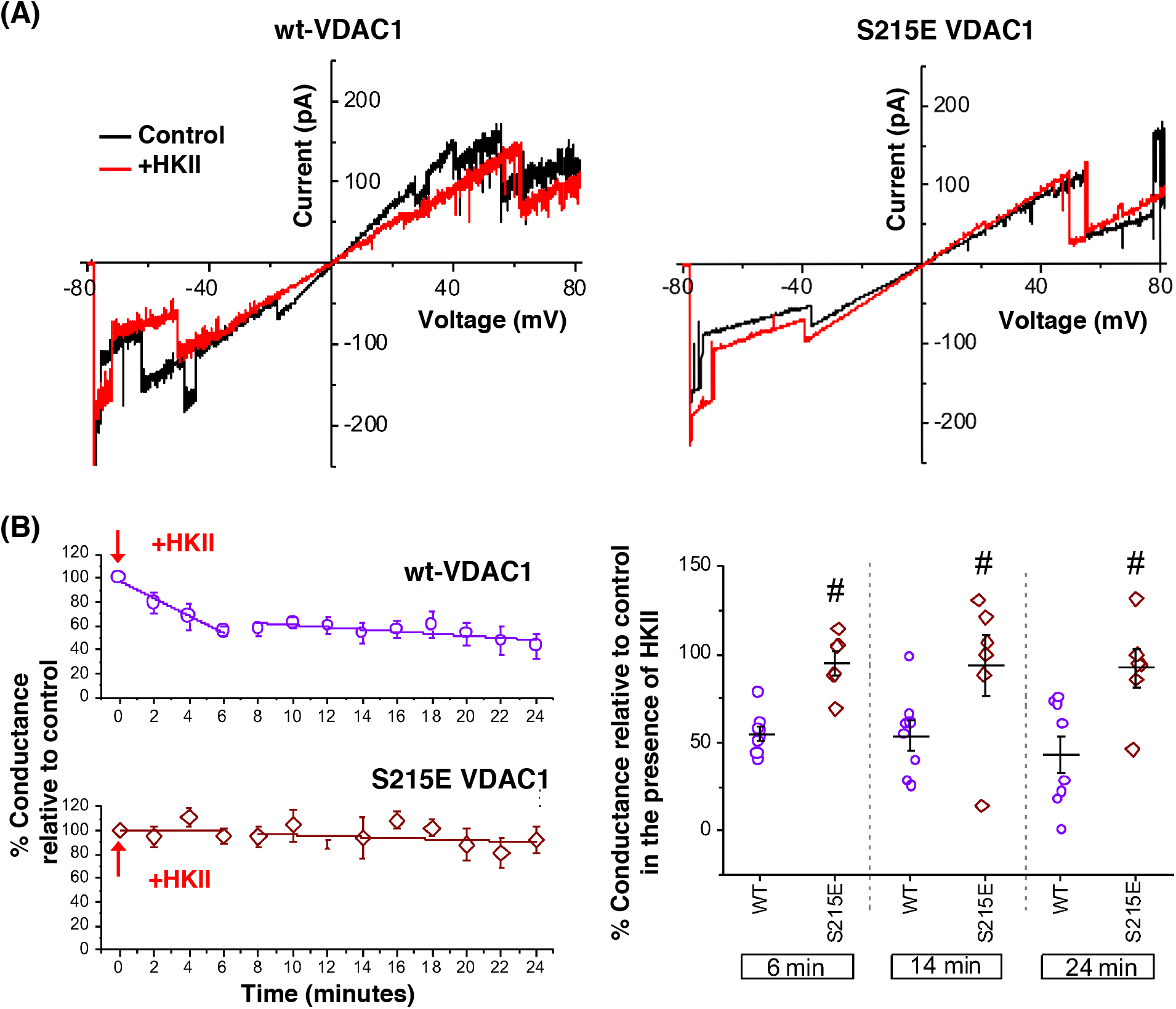
Effects of HKII on wt-VDAC1 and S215E phosphomimetic mutant. (A) Current-voltage relationships for wt-VDAC1 (left) and S215E VDAC1 (right) in the absence (control; black lines) or the presence of HKII (+HKII; red lines), acquired during a −80 to +80 mV linear voltage ramp protocol. (B) Effects of HKII on VDAC1 conductance monitored over time (purple circles: wt-VDAC1; brown diamonds: S215E phosphomimetic mutant). Change in conductance in the presence of HKII was relative to that prior to the addition of HKII. Summary graphs are shown for both wt-VDAC1 (*n* = 8) and S215E VDAC1 (*n* = 6) in the presence of HKII. Representative steady-state current recordings for both wt-VDAC1 and S215E VDAC1 in the presence and absence of HKII are shown in Fig. S5.

### Phosphorylation of VDAC1 disrupts HKII binding

The immense HKII/VDAC1 interaction energy shown in MD simulations (Fig. S2B) and the composition of the HKII/VDAC1 binding interface (Fig. S3A) strongly suggests the binding interactions between HKII and VDAC1 are dominated by salt bridges. Therefore, their complex formation might be interfered or disrupted with the presence of additional charged species, such as post-translational modifications to the proteins. Coincidently, an earlier proteomics screening of phosphorylated mitochondiral proteins have identifed S215 of VDAC as an phosphorylation site *in vivo*^52^, which is locating at the HKII/VDAC1 binding interface (Fig. S3A).

To investigate the impact of S215 phosphorylation on complex formation between HKII and VDAC1, we repeated the BD simulations after introducing a phosphoserine residue at position 215. Whereas BD simulation of wt-VDAC1 resulted in an HKII-binding hot spot near S215 (Cluster 1), the probability of HKII presence at this region is substantially reduced after S215 phosphorylation (Fig. 6A), suggesting that the formation of HKII/VDAC1 complex can be inhibited when S215 is phosphorylated. The dissappearing of population corresponding to Cluster 1 can also be predicted by the greatly reduced HKII/VDAC1 interaction energy if a phosphoserine is directly placed at S215 position of the BD trajectories of wt-VDAC1: the electrostatic interactions between the two proteins suffered from ~2.6-fold reduction on average in Cluster 1, while other clusters showed only minor changes (Fig. S6).

**Figure 6.**
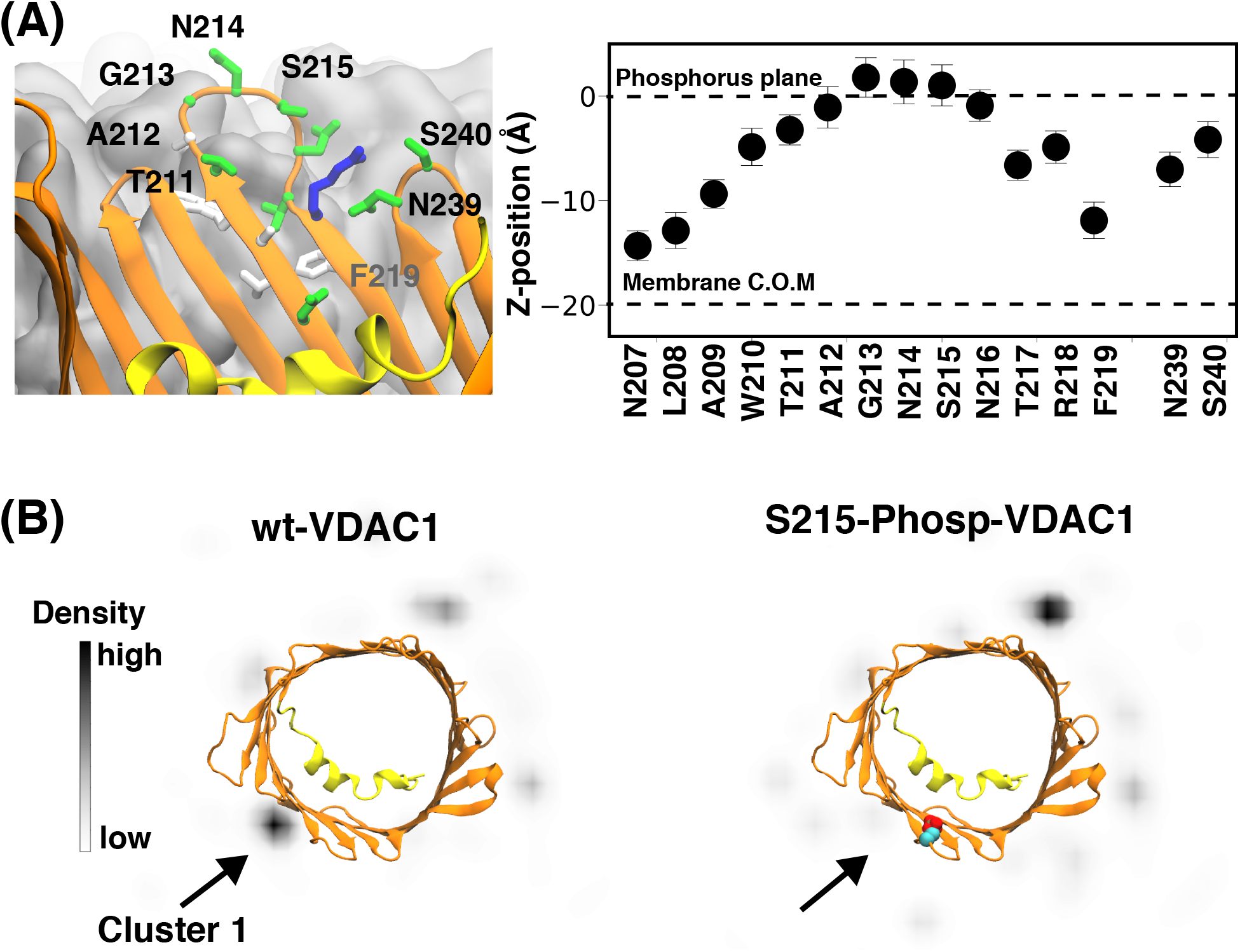
Phosphorylation of S215 in VDAC1 disrupts HKII/VDAC1 binding. (A) Top-down (cytosolic) view of the *xy* projection of densities of the C.O.M of the H-anchor bound to VDAC1 during the BD simulations. The densities are obtained using all 100 independent replicates of the BD simulations for wt-VDAC1 (left) and S215-phosphorylated (S215-Phosp-VDAC1, right). (B, *Left*) Cytosolic accessibility of S215. VDAC1 barrel viewed from the membrane plane, highlighting the position of S215 at a location accessible to bulk solvent. The membrane is colored in gray. (B, *Right*) Mean and standard deviation of the relative positions along the membrane normal (*z*-position) of the C.O.M the side-chain of highlighted VDAC1 residues (*light*), derived from a 200-ns MD simulation of membrane-embedded VDAC1.

VDAC1 phosphorylation is known to modulate HKII binding *in vivo*^42^. Therefore, the impact of VDAC1 S215 phosphorylation to HKII/VDAC1 interactions was further corroborated with electrophysiology experiment using a phosphomimetic mutant of VDAC1, S215E. Comparing to wt-VDAC, the HKII-induced partial blockage is rarely observed, if not completely abolished, in membranes containing the S215E phosphomimetic mutant of VDAC1 (Fig. 5B, S5). Additionally, the presence of HKII does not alter the current-voltage characteristics of S215E-VDAC1 significantly (Fig. 5A). Based on these observations, the S215E point mutation appeared to prevent or diminish the interaction of HKII with VDAC1, supporting the predicted outcome of the computational model.

To date, it remains unclear which kinase or signaling pathway mediates VDAC1’s phosphorylation at S215 position. We used the peptide sequence near S215 for kinase substrate prediction and did not yield a convincing kinase hit. However, since HKII/VDAC1 binding can be inhibited by the S215 phosphorylation, the kinase responsible for S215 phosphorylation is expected to belong to a pro-apoptotic pathway or at least not in a pathway promoting cell survival. Intriguingly, VDAC1 is a known substrate of glycogen synthase kinase (GSK)-3*β* ^53^, a major promoter of mitochondrial intrinsic apoptotic pathway^54^. VDAC phosphorylation by GSK3*β* is also linked to a reduced HKII binding^53^, and the disruption of HK binding to VDAC1 has been shown to play an essential role in several diseases including cardiac ischemia-reperfusion injury^55–57^, by promoting mitochondria-mediated cell death^7,53,58^. The matching phenotype leads us to speculate that the kinase responsible for VDAC1 S215 phosphorylation might be GSK3*β*.

GSK3*β* is known to highly favor a “primed” substrate where another phosphorylated serine/threonine is locating at the +4 residue position downstream of the phosphorylation site^59^. One may argue that in VDAC1, such position is a membrane-embedded F219, which cannot serve as a priming residue for GSK3*β* recognition (Fig. 6B). Based on crystal structures of GSK3*β* with a bound pseudo-substrate phosphopeptide inhibitor^60^, GSK3*β* seems to recognize substrate peptides with an extended backbone configuration where the priming phosphate sits ~13 Å away from the transferring *γ*-phosphate. Interestingly, the *β*-barrel VDAC is full of extended backbones and its cytoplasmic rim has several serine and threonine residues, among them S215 is locating at a *β* -turn slightly protruding out of the membrane surface and is exposed to the cytosol (Fig. 6B). In addition, residue S240 is locating at the immediately following *β*-turn at the cytoplasmic side, and is in line with the downstream residues of S215, spacing on average 14.3±1.7 Å between the two hydroxyl groups in our simulations (Fig.6B). Most importantly, S240 was also found phosphorylated *in vivo* in the mitochondrial proteomics study^52^. The proximity of phospho-S240 from S215 and the extended backbone configuration of residues between them make it possible for GSK3*β* to bind to the cytoplasmic rim of VDAC1, recognizing phospho-S240 and phosphorylating S215.

Although the consensus sequence of GSK3*β* substrate peptides S/T-X-X-X-S/T(P)^61^ would predict a subsequent T211 phosphorylation if S215 is phosphorylated by GSK3*β*, the two residues are situated right at the two ends of the same *β* -turn (Fig. 6B). It is therefore unlikely that phospho-S215 can be the primer for GSK3*β* to phosphorylate T211 of VDAC1 due to incompatible backbone configuration. To date, we have yet to discover a literature reporting T211 phosphorylation of VDAC.

## Concluding Remarks

Atomic structures of protein complexes are usually derived from experimental techniques such as X-ray crystallography and cryo-EM. In parallel, and particularly for cases such as membrane-bound systems where obtaining detailed structures might be too challenging, computational techniques such as molecular docking, BD or MD simulations offer powerful complementary tools^62,63^. Here, using a hybrid computational approach combining MD and BD simulations we report a stable structural model for the HKII/VDAC1 complex, which we validate using complementary experiments at multiple levels. The complex partially blocks the permeation pathways of the VDAC1 channel, a key feature that was also observed in our electrophysiology measurement, strongly supporting the relevance of the derived complex structure. Furthermore, the binding interface of the complex is highlighted by the presence of a potential phosphorylation site S215, whose phosphorylation/phosphomimetic-mutation results in inhibition of HKII/VDAC1 interactions both in our BD simulations and in our electrophysiology recordings. This inhibition appears to be mediated through disruption of the electrostatic attraction between H-anchor and cytosolic outer-rim of VDAC1.

Though other phosphorylation sites on VDAC1 have been identified^2,56^ and several of these sites have been implicated in cardiac and neurodegenerative diseases, their functional consequences have not been delineated. Our study shows that phosphorylation of a single residue, S215, can disrupt HKII interaction with VDAC1. This provides insight into how phosphorylation of specific sites on VDAC1 can impact its binding with HKII and potentially other proteins that translocate to mitochondria, for example the pro-apoptotic protein BAX. Characterizing the complex molecular interactions between VDAC1 and anti- and pro-apoptotic proteins will help delineate their roles in cell death and survival.

## Methods

### Membrane binding simulations

We used the X-ray structure of human HKII (PDB ID: 2NZT) as a starting point for our simulations. Glucose and glucose-6-phosphate were removed from the crystal structure, and one monomer from the HKII dimer was selected for the simulations. The missing regions (residues 1-16, 98-104, 346, 404-405, 518-525, 546-552, 645-649, 914-917) from the crystal structure were modeled using MODELLER^64^. Among missing regions, residues 1-16 were predicted to be in an *α*-helical structure by the secondary structure prediction tool YASPIN^65^, while other missing regions were predicted to be unstructured. As residues 1-16 are a part of the H-anchor and known to be crucial for membrane-binding, we carefully modeled this part by using as a template the highly homologous isoform enzyme from rat, HKI (PDB ID: 1BG3), for which the H-anchor is fully resolved as an *α*-helix. Notably, the H-anchor of rat HKI shares high sequence similarity with human HKII (Fig. S1). Other missing regions were modeled as loops using conjugate gradients and molecular dynamics with simulated annealing approach in MODELLER^66^.

To reduce the system size in our membrane-binding simulations, we removed the C-domain and used only the N-domain (residues 1-463) of human HKII for membrane binding simulations. N-domain of human HKII, referred to as HKII-N hereafter, is known to be sufficient for binding to the outer membrane of mitochondria (OMM)^45^. In order to assign the protonation states of ionizable residues of HKII-N, pK_*a*_ values were estimated using PROPKA^67,68^. Then, HKII-N was placed above a symmetric HMMM lipid bilayer composed of 108 phosphatidylcholine (PC), 58 phosphoethanolamine (PE), 26 phosphatidylinositol (PI), 4 phosphatidylserine (PS) and 2 phosphatidic acid (PA) lipid molecules in each leaflet. The lipid composition was chosen to approximately replicate the OMM of mammalian cells^69^. HKII-N was placed 5 Å above the membrane, where 5 Å is the minimum distance between any HKII-N atom and any membrane atom along the membrane normal (z-axis). Ten independent HMMM systems were constructed using CHARMM-GUI^70^, each resulting in a box with dimensions of 125 125 174 Å^3^, including 240,000 atoms. To minimize possible bias in membrane binding simulations due to a particular initial lipid distribution, the lipid bilayers in all replicas were generated independently using CHARMM-GUI^70^ to obtain a different (randomized) initial lipid distribution for each replica. All replicas were neutralized with 150 mM NaCl and solvated with the TIP3 water^71^. All the 10 solvated HMMM systems were then energy-minimized for 2,500 steps before used for the subsequent 200 ns membrane binding simulations. A harmonic restraint on the z-position (along the membrane normal) was applied to the carbonyl atoms of short-tailed lipids with a force constant of *k* = 0.1 kcal mol^−1^ Å^−2^, to mimic the atomic distributions of a full-tailed lipid bilayer and to prevent occasional diffusion into the aqueous phase^72,73^.

### Conversion of HMMM lipids to full-tailed lipids

In order to generate a complete model of membrane-bound HKII-N, we used the last frame of one of the HMMM membrane-binding simulations described above and converted it to a full-membrane construct using CHARMM-GUI^70^. Short-tailed HMMM lipids were transformed into full lipids by removing the DCLE molecules and adding the missing carbons to the lipid tails while preserving the positions of the headgroups and initial six carbons of the lipid tails. The system was then energy-minimized for 10,000 steps and then equilibrated for 2 ns, while harmonic restraints (force constant, *k* = 1 kcal mol^−1^ Å^−2^) were applied to (1) the heavy atoms of the protein and (2) the positions of lipid heavy atoms corresponding to short-tailed lipids only in the *cis* leaflet of the lipid bilayer (the leaflet facing the protein). These restraints were applied to preserve lipid-protein interactions established during the association of the protein with the lipid membrane in HMMM simulations, while the newly added carbon atoms adjusted to the system^74^. Following this step, the system was simulated without restraints for 200 ns, resulting in a fully relaxed, membrane-bound protein.

### Simulation of membrane-embedded VDAC1

The X-ray structure of mouse VDAC1 (PDB ID: 3EMN)^40^ was embedded in a full membrane, with the same lipid composition as described above, using CHARMM-GUI^31^. Mouse and human VDAC1 proteins have a sequence similarity of 94%. The system was neutralized with 150 mM NaCl and solvated with TIP3 water^71^ resulting in a box size of 120 × 120 × 84Å^3^ including 110,000 atoms. Then the system was energy-minimized for 2,500 steps and equilibrated for 1 ns, during which C*α* atoms of the protein were harmonically restrained with a force constant, *k*, of 1 kcal mol^−1^ Å^−2^. Following this step, the system was simulated without restraints for 200 ns.

### MD simulation of HKII/VDAC1 complexes

For each of the complexes (HKV1, HKV2 and HKV3) derived from the BD simulations, a full-length HKII/VDAC1 model was generated by adding the C-domain of HKII to the coordinates of the HKII-N/VDAC1 complex (based on the known full structure of HKII). Each extension was carried out by adopting coordinates from full-length HKII after superimposing the backbone atoms of residues S449-L463 (an *α*-helix) into HKII-N. The resulting full-length models were then embedded in a full-membrane with the same lipid composition as described above using CHARMM-GUI^31^. These three different systems were neutralized with 150 mM NaCl and solvated with TIP3 water^71^. HKV1 simulation system resulted in a box size of 215 × 215 × 160 Å^3^ including 700,000 atoms; HKV2 system resulted in a box size of 120 × 120 × 240 Å^3^ including 330,000 atoms; and HKV3 system resulted in a box size of 160 180 140 Å including 330,000 atoms. Each system was energy minimized for 2,500 steps and equilibrated for 1 ns while harmonically restraining the C*α* atoms of both proteins with a force constant, *k*, of 1 kcal mol^−1^ Å^−2^. Then, HKV1 and HKV3 systems were simulated for 350 ns, while HKV2 was simulated for 80 ns, without restraints. HKV2 was simulated shorter since it exhibited separation of the two proteins early on (starting at ~30 ns; Fig. S2), and therefore longer simulations were not pursued.

### MD simulation protocols

MD simulations in this study were performed using NAMD 2.13^75,76^ utilizing CHARMM36m^77^ and CHARMM36^78^ force field parameters for proteins and lipids, respectively. Bonded and short-range nonbonded interactions were calculated every 2 fs, and periodic boundary conditions were employed in all three dimensions. The particle mesh Ewald (PME) method^79^ was used to calculate long-range electrostatic interactions every 4 fs with a grid density of 1 Å^−3^. A force-based smoothing function was employed for pairwise nonbonded interactions at a distance of 10 Å with a cutoff of 12 Å. Pairs of atoms whose interactions were evaluated were searched and updated every 20 fs. A cutoff (13.5 Å) slightly longer than the nonbonded cutoff was applied to search for the interacting atom pairs. Constant pressure was maintained at a target of 1 atm using the Nosé-Hoover Langevin piston method^80,81^. Langevin dynamics maintained a constant temperature of 310 K with a damping coefficient, *γ*, of 0.5 ps^−1^ applied to all atoms. HMMM simulations were performed by employing a constant area in the XY dimension (membrane plane). For the full-membrane simulations, a constant ratio was used instead, which keeps the X:Y ratio of the unit cell constant. Simulation trajectories were collected every 10 ps.

### Electric field simulations

Ionic current through VDAC1 was calculated by performing simulations of the membrane-embedded form of the channel in the presence of a constant electric field normal to the membrane. Simulations were performed for both HKII-free and HKII-bound states (HKV1) of VDAC1. The starting point for each simulation was the last snapshot of their respective full-membrane equilibrium MD simulation. For each system, the salt concentration was increased from 150 mM to 500 mM by using the AUTOIONIZE plugin of VMD^82^, in order to enhance the number of ion permeation events during the electric field simulations. Three independent simulations were performed for each system, under a membrane potential difference of −50 mV, each for 80 ns.

Ionic current (I) was computed by counting the number of ions (Na^+^ and Cl^-^) that cross the VDAC1 channel (moving all the way from one side of the membrane to the other) over time, i.e. 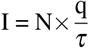, where N is the number of crossing events over a time interval *τ*, and q is the charge of the ion (1.60217662×10^−19^ Coulombs for Na^+^ and −1.60217662×10^−19^ Coulombs for Cl^−^).

### Brownian dynamics (BD) simulation setup

We used the membrane-bound HKII-N and membrane-embedded VDAC1 to simulate the formation of their complexes using BD simulations. Atomic coordinates of both proteins were extracted from the final snapshot of their respective MD simulations in full-membranes. During the BD simulations, membrane-bound HKII-N was considered as the moving protein, while VDAC1 was the stationary protein, with both proteins modeled as rigid body entities.

Given the rigid representation of the molecular entities in BD simulations, we did not use an explicit membrane in these simulations. In order to take into account membrane placement of HKII-N and VDAC1 (as observed in the MD simulations of the membrane-bound forms of these proteins), in the beginning of each simulation, the z-coordinates of each protein were shifted so that their respective membrane midplanes would match (Fig. 3A). Another set of restraints were then used to maintain both the insertion position and orientation of HKII-N during the BD simulation; we applied restraints on the positions of H-anchor residues along the z-axis in our BD simulations. Restraints were applied on four residues (MET1, PHE10, ASN15 and GLN20) of H-anchor using a grid-based potential coupled to each residue. Grid-based potentials were obtained by Boltzmann inversion of the probability distribution function of the z-coordinate of each residue obtained from the last 100 ns of membrane-bound MD simulation of HKII-N in full-membranes. Applied restraints were able to maintain the positioning and the tilt angle distribution of the H-anchor in BD simulations close to those observed in full-membrane MD simulations (Fig. 3A), while allowing for the lateral diffusion of HKII-N around VDAC1 and their complex formation. To avoid unnecessary sampling of HKII-N during simulation, we employed a circular grid-based potential wall around VDAC1 with a radius of 150 Å.

### BD simulation protocols

BD simulations were performed using a GPU accelerated BD code, ATOMIC RESOLUTION BROWNIAN DYNAMICS (ARBD)^83^. The masses and moments of inertia of the HKII-N (moving protein) were calculated directly from its atomic coordinates. HYDROPRO program suite^84^ was used to estimate the translational and rotational friction coefficients of HKII-N protein which provided Langevin forces and torques at each timestep to keep the system at 310 K. To be noted, our calculated friction coefficients represent the diffusive behavior of HKII-N in aqueous solutions. Though this diffusive property would be significantly different for membrane-bound HKII-N, it will not affect the results of our study, as we are not focusing on either diffusive or kinetic properties of HKII-N binding to VDAC1.

The force and torque acting on HKII-N, due to stationary protein VDAC1, were calculated as follows. First, a grid of electric charge and Lennard-Jones (LJ) particle densities was obtained from the atomic coordinates of membrane-bound HKII-N with a 1 Å resolution. The density in each grid cell experienced a local force due to the corresponding grid-specified stationary VDAC1 potential (electrostatic or LJ terms). These local forces and the corresponding torques were summed over to obtain the total force and torque on the HKII-N molecule. A cutoff of 34 Å was used for the force calculation. Electric charge and LJ particle density of the atoms comprising HKII-N were calculated using the VOLMAP utility in VMD. On the other hand, stationary protein VDAC1 was represented by the electrostatic and LJ potential map of its atomic coordinates. The electrostatic potential map for the stationary protein (VDAC1) was calculated by solving the nonlinear Poisson Boltzmann equation implemented in the APBS software^85,86^. *cglen* and *fglen* options in APBS were chosen to ensure a resolution of 1 Å for computing the cubic electrostatic potential grids. Dielectric constants for the protein interior and solvent were set to 12.0 and 78.0, respectively. Ionic radii were set as per the CHARMM36 force field^77^. A cubic-spline surface^32^ model was implemented to model the dielectric interface and ion accessibility. The rate of dielectric transition was set to 0.3 and surface density to 10. Radius of the solvent molecule was set to 1.4 Å, same as water. These parameters are comparable to previous studies of protein-protein interactions using ARBD simulations^87^.

The LJ parameters of the atoms comprising the stationary protein (VDAC1) were clustered into three categories: one representing all hydrogen atoms, another representing oxygen and nitrogen atoms, and the final one representing carbon and sulfur atoms. The atoms in each category were assigned an average value for the parameters R_min_ and *ε*. Then, *ε* was scaled down by 0.3, following a similar approach adopted by McGuffee et al.^88^ to avoid stickiness of proteins during the BD simulations. These two parameters, R_*min*_ and scaled *ε*, were used to generate potential maps for the interaction of the three above-mentioned atom categories with the stationary protein, VDAC1, using the implicit ligand sampling (ILS) implementation of VMD at 1 Å resolution^82,89^. ARBD simulations were performed using a timestep of 200 fs. Simulation coordinates were collected every 0.2 nanosecond.

### Generation of an H-anchor plugged model

An alternative model for the HK/VDAC1 complex has been reported in the literature^38,39^. In this model, which we refer to as the “H-anchor plugged model”, the H-anchor completely plugs the VDAC1 pore. To investigate the electrophysiological properties of such a model and how VDAC1 permeability is affected, we generated an approximate model of H-anchor plugged HKII/VDAC1 complex by docking the H-anchor into the lumen of VDAC1, which was then used to in our electric field simulation for current measurements. Docking was performed using the “easy interface” implemented in HADDOCK2.2 web portal^90^. The X-ray structure of human HKII (PDB ID: 2NZT, missing regions modeled) and the same of mouse VDAC1 (PDB ID: 3EMN)^40^ were used for doking. Docking was performed between the residues of H-anchor and the residues of VDAC1 facing the lumen on the cytoplasmic half resulting in multiple structural models. A representative structure was selected based on the best HADDOCK score, which was then embedded in a full-membrane with the same lipid composition as those used for our own model (described above) using CHARMM-GUI^31^. The system was neutralized with 150 mM of NaCl and solvated with TIP3 water^71^, resulting in a box size of 120 120 200 Å^3^ including 270,000 atoms. The newly generated system was energy minimized for 2,500 steps and equilibrated for 1 ns while harmonically restraining the *Cα* atoms of both of proteins with a force constant of *k* = 1 kcal mol^−1^ Å^−2^. Then, the system was simulated without restraints for 100 ns before using it for computational electrophysiological measurements.

### Analysis

System visualization and analysis were carried out using VMD^82^. Depth of insertion of HKII-N into the membrane was assessed by calculating the z component of the center of mass (C.O.M) of protein side-chain atoms relative to the average plane of the phosphorous atoms of the membrane. Interaction energy (van der Waals + electrostatic) between HKII-N and VDAC1 was calculated using the NAMDENERGY plugin of VMD. Contacts between HKII and VDAC1 are calculated using a 3 Å distance cutoff between any atoms of the two proteins. A hydrogen bond was counted to be formed between an electronegative atom with a hydrogen atom (H) covalently bound to it (the donor, D), and another electronegative atom (the acceptor, A), provided that the distance D-A is less than 3 Å and the angle D-H-A is more than 120^◦^. Clustering was performed based on the position of the HKII-N with respect to VDAC1 (calculated using the root-mean-square deviation (RMSD) of HKII-N after superpositioning the BD trajectories using VDAC1). The “measure cluster” module implemented in VMD^82,91^ was used for clustering analysis with an RMSD cutoff of 10 Å.

### Generation of VDAC1 mutant construct

Full-length cDNA of rat VDAC1 (GenBank: BC072484, purchased from Open Biosystems) was first subcloned into pET21a vector (Novagen) to generate pET-VDAC1 (wt). The VDAC1 mutations at serine 215 to glutamate (S215E) was generated by PCR with QuikChange site-directed mutagenesis kit (Agilent Technology) using pET-VDAC1 as cDNA templates and the following primers: S215A: Forward 5’-CTC GCC TGG ACC GCA GGA AAC GCT AAC ACT CGC TTT GG-3’ and Reverse 5-‘ CC AAA GCG AGT GTT AGC GTT TCC TGC GGT CCA GGC GAG-3’ ; S215E Forward: 5’-CTC GCC TGG ACC GCA GGA AAC GAG AAC ACT CGC TTT GG-3’ and Reverse 5-‘CC AAA GCG AGT GTT CTC GTT TCC TGC GGT CCA GGC GAG -3’. The plasmids pET-VDAC1, pET-VDAC1-S215A and pET-VDAC1-S215E were separately transformed into BL21 *E. coli* for protein expression. Cells were grown at 37°C in LB medium to A600= 0.6 and VDAC1 protein expression was induced with 1 mM IPTG for overnight. After induction, VDAC1 proteins were extracted with BugBuster Master Mix (Novagen) and purified with Ni-NTA His·Bind Resins (Novagen). Purified VDAC1 was refolded at 4° by slowly dropwise dilution of one volume protein into 10 volumes refolding buffer containing 20 mM Tris-Cl, 100 mM NaCl, 1% lauryldimethylamineoxide (LDAO), and 1 mM DTT at pH 7.4 with slowly stirring. The refolded VDAC1 protein was dialyzed against modified refolding buffer (0.1% LDAO and without DTT) at 4°C to decrease LDAO concentration and to remove DTT.

### Experimental electrophysiology

Recombinant rat wild-type (wt) or mutant (S215E) VDAC1 proteins were reconstituted into planar lipid bilayers as described previously^51^ with some modifications. Briefly, phosphatidylethanolamine (PE) and phosphatidylcholine (PC) (Avanti Polar Lipids) were mixed in a ratio of 7:3 (v/v), dried under nitrogen gas, and resuspended in n-decane (Sigma) for a final lipid concentration of 25 mg/ml. The *cis*/*trans* chambers contained symmetrical solutions of 10 mM Trizma base (Sigma), 500 mM KCl (Sigma) and 1 mM CaCl2 (Sigma), pH 7.4. The *cis* chamber was held at virtual ground and the *trans* chamber was held at the command voltage. The pClamp software (version 10, Molecular Devices, San Jose, CA) was used for data acquisition. Currents were digitized at 5 kHz and low pass filtered at 1 kHz using a voltage clamp amplifier (Axopatch 200B, Molecular Devices) via a digitizer (DigiData 1440A, Molecular Devices). The recombinant VDAC1 proteins were added into the *cis* chamber. Insertion of VDAC1 into the bilayer membrane and its function were monitored and confirmed by current recordings in response to a ramp protocol from 80 to +80 mV, as VDAC1 is characterized by its uniquely distinct current-voltage relationship. Subsequently, the solution in the *cis* chamber was replaced with the same initial buffer solution at the speed of 2.5 ml/min to remove non-inserted VDAC1 proteins and prevent additional VDAC1 insertion into the bilayer membrane. Current recordings under control conditions (in the absence of HKII) were initially taken, followed by addition of HKII (human recombinant HKII, 60 kU/ml; Genway Biotech, Slan Diego, CA) into the *cis* chamber. Currents were monitored during a 30-s recording duration every two minutes up to 24 minutes, and analyzed using Clampfit (Molecular Devices) and Origin (version 10; OriginLab, Northampton, MA). Mean current from each time point was normalized to its control as a percentage data, which was used for statistics later.

Significant differences between groups were determined with one-way ANOVA (SPSS Statistics 24, IBM) with post-hoc significance analysis. P < 0.05 was considered significantly different.

## Acknowledgements

We thank Christopher Maffeo (UIUC) for insightful conversations and assistance designing Brownian dynamics simulations. This research was supported by the National Institutes of Health grants R01-HL131673 (to WMK, AKSC and ET), and P41-GM104601 (to ET). Simulations in this study have been performed using allocations at National Science Foundation Supercomputing Centers (XSEDE grant number MCA06N060) and the Blue Waters supercomputer of National Center for Supercomputing Applications at University of Illinois at Urbana-Champaign.

## Author contributions statement

N.H., P.W., A.C., W.K., and E.T. designed research; N.H., Q.C., M.Y., and G.N. performed research; N.H., P.W., and W.K. analyzed data; and N.H., P.W., W.K., and E.T. wrote the paper.

