## Supplementary Table and Figures for "Structural Basis of Complex Formation Between Mitochondrial Anion Channel VDAC1 and Hexokinase-II"

### Supporting Information

**Table S1.** Population and number of contacts between H-anchor and VDAC1 for five clusters.

| Cluster | Population (%) | H-anchor contacts |
|---|---|---|
| 1 | 14.0 | 13.3±6.8 |
| 2 | 9.3 | 3.9±2.2 |
| 3 | 6.0 | 8.4±5.7 |
| 4 | 4.8 | 0.0±0.0 |
| 5 | 4.6 | 0.0±0.2 |

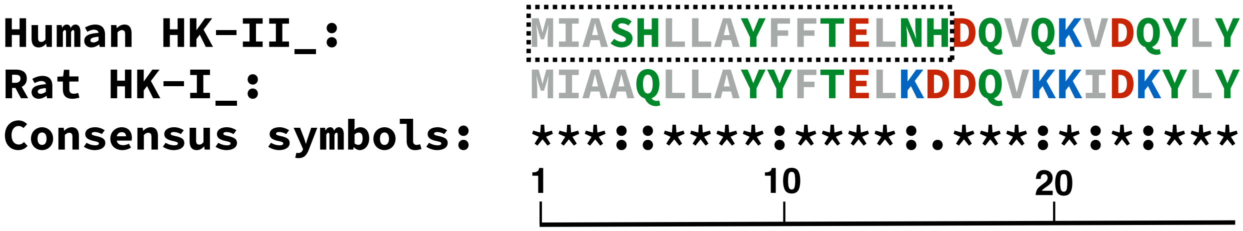


**Figure S1.** Sequence alignment of H-anchor of human HKII and rat HK-I. Residues from 1 to 16 of human HKII (highlight with dotted box) is missing from the crystal structure (PDB ID: 2NZT). Description of the consensus symbols: An * (asterisk) indicates positions that have a fully conserved residue, A : (colon) indicates conservation between residues of strongly similar properties, A . (period) indicates conservation between residues of weakly similar properties. Each residue is represented by their one-letter amino acid abbreviation and colored based on their residue type: Gray representing hydrophobic, green polar, red acidic, and blue basic residues. The alignment was performed on full-length human HKII and full-length rat HK-I using the Clustal Omega[92] alignment tool within Uniprot[93]. Full-length human HKII and rat HK-I share an overall 73% sequence identity.

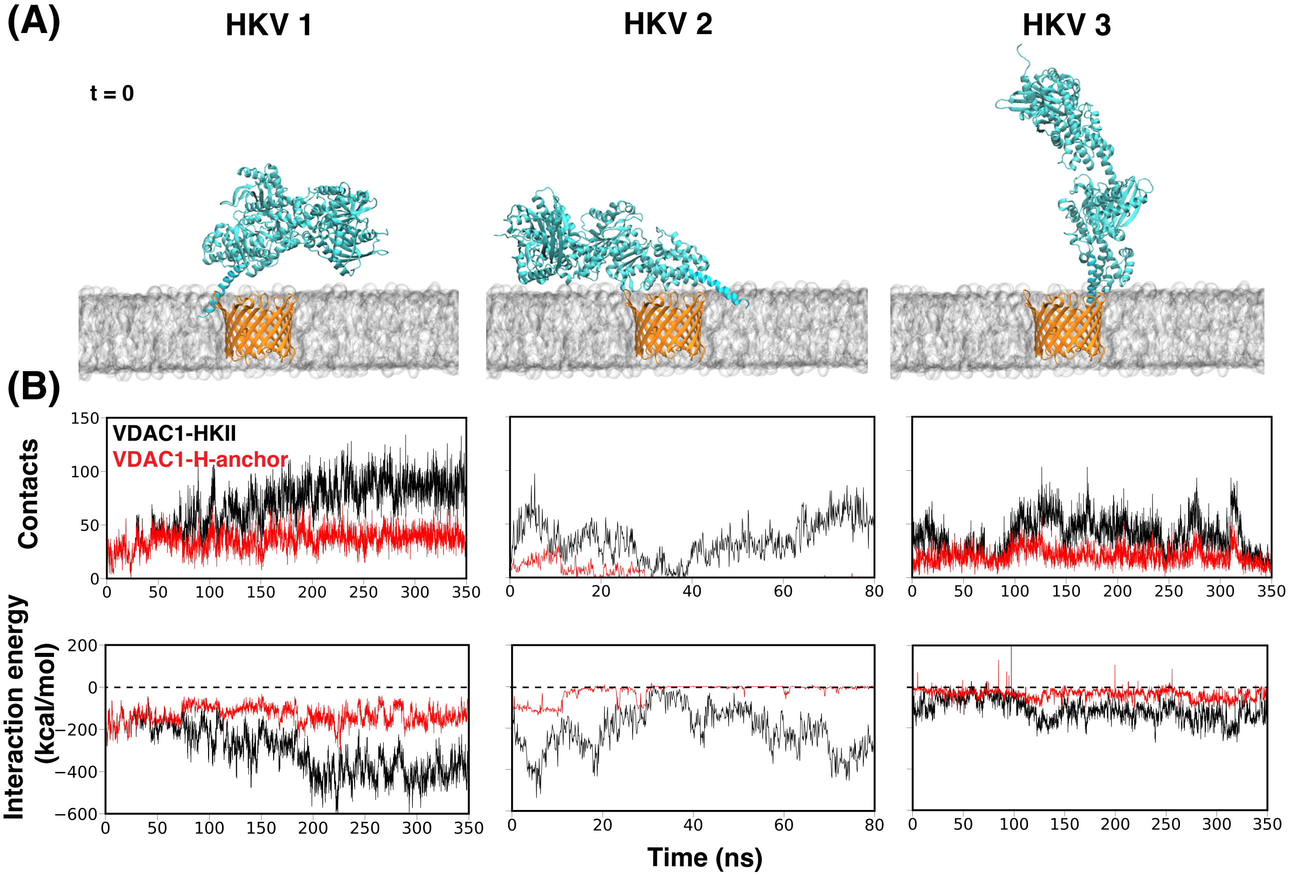


**Figure S2.** (A) Molecular representation of membrane-embedded HKV1, HKV2 and HKV3 at time = 0. (B) Time evolution of the number of contacts and interaction energy (van der Waal + electrostatic) between HKII and VDAC1 during MD simulation of the three complexes.

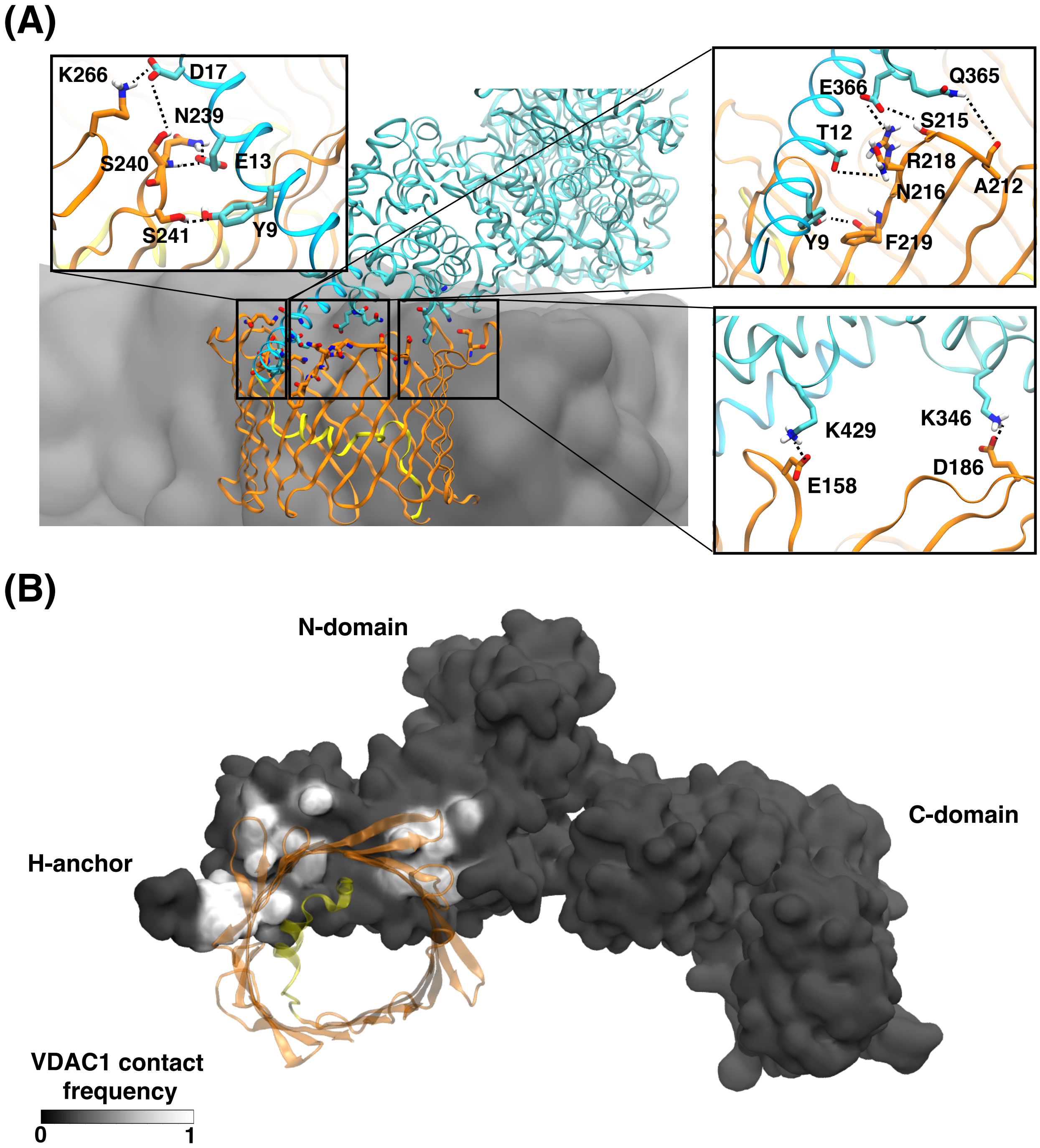


**Figure S3.** (A) Hydrogen bond and salt-bridge interactions between residues of VDAC1 and HKII maintaining the HKV1. Interactions that have a significant presence (>10% occurring probability) in the simulation are shown in the *Inset*. The membrane is shown in gray. (B) VDAC1 contact frequencies mapped onto the surface of HKII (black and white), viewed from the mitochondrial side and perpendicular to the membrane surface. The calculation was performed using the last 150 ns of MD simulation during which HKII/VDAC1 contact converges. VDAC1 is shown in transparent representation, colored in orange.

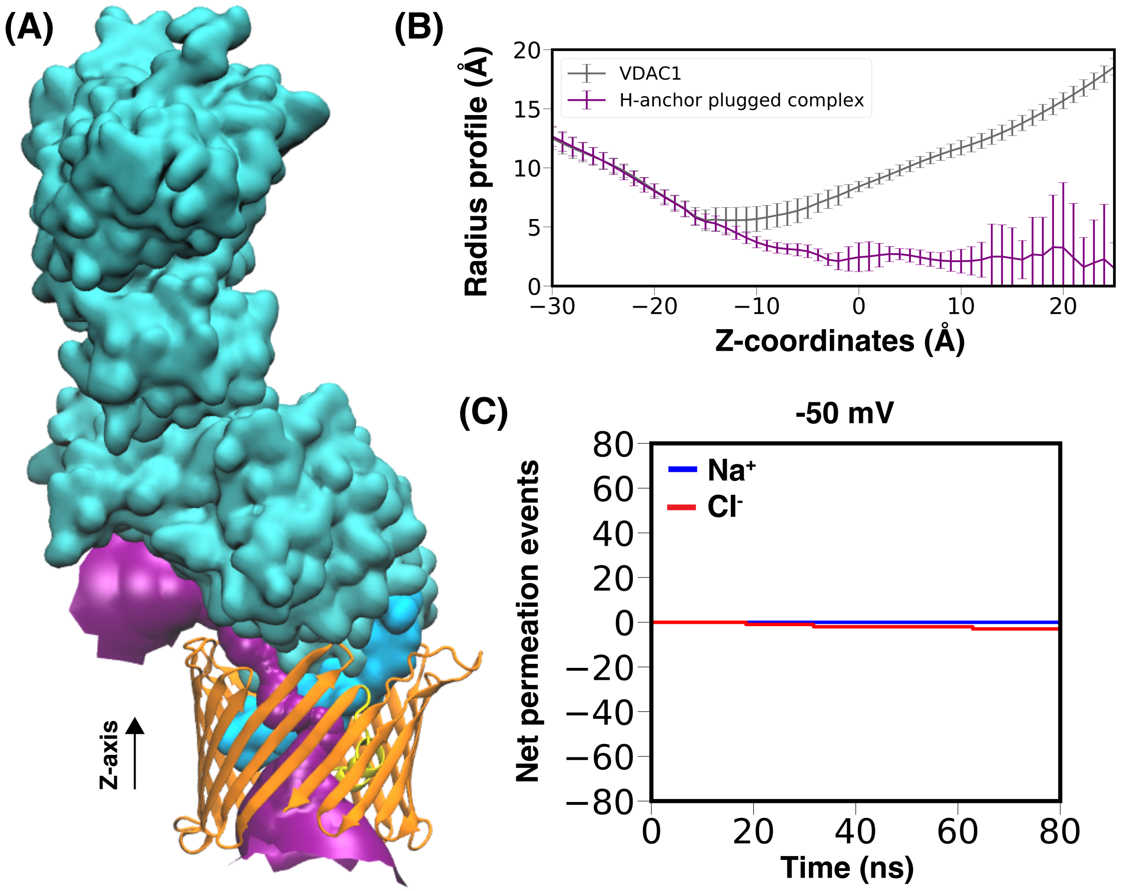


**Figure S4.** HKII/VDAC1 interaction in a H-anchor plugged model. (A) Purple surface showing VDAC1 pore radius profile in HKV1, made with the program HOLE[50]. Z-axis represents the membrane normal. (B) The average radius of the VDAC1 pore calculated using HOLE[50]. (C) The cumulative net number of channel-crossing events by $Cl^-$ (red traces) and $Na^+$ (blue traces), tracked over the time course of the electric field MD simulation at −50 mV for the H-anchor plugged complex.

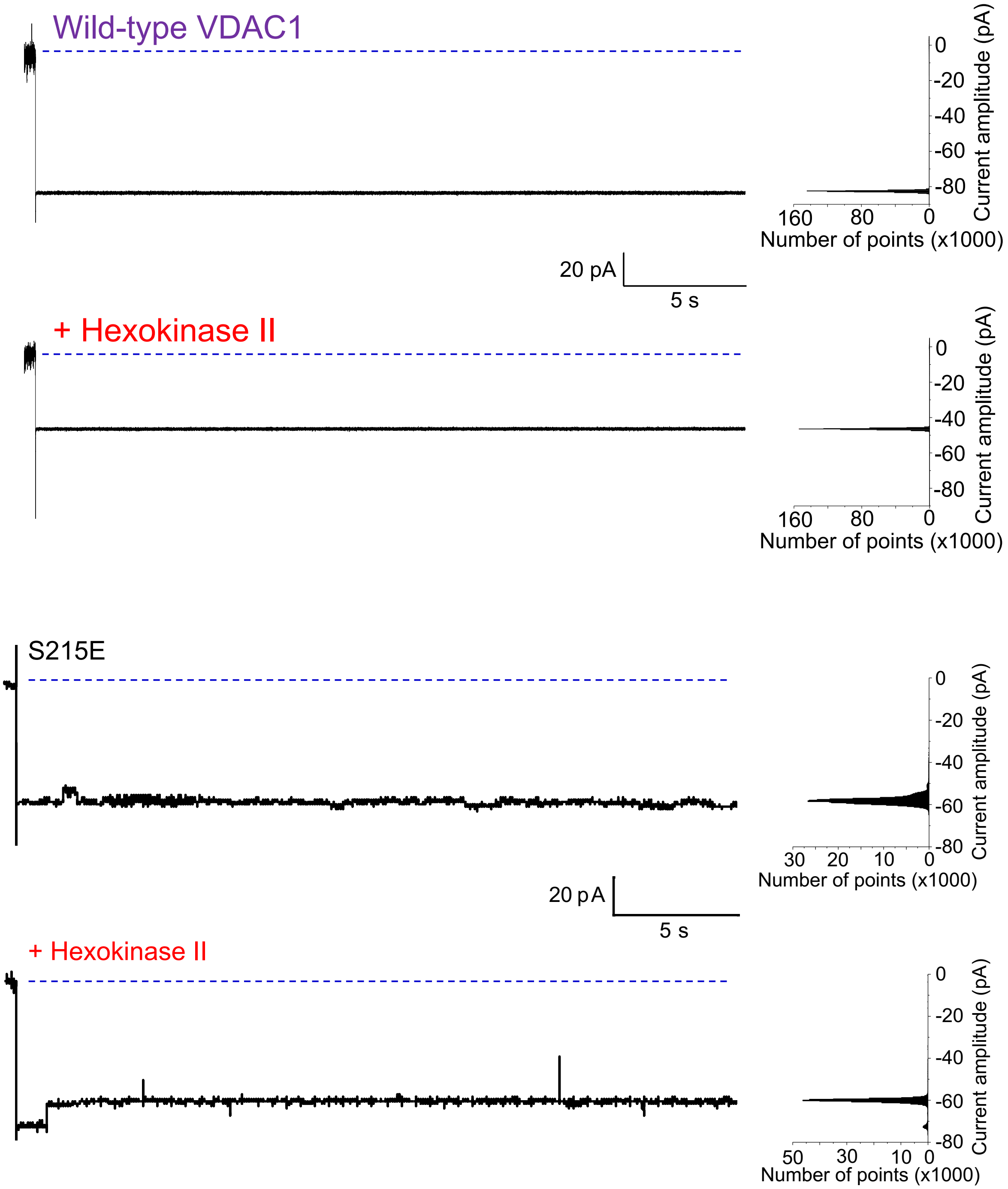


**Figure S5.** Effect of HKII on the conduction of wt-VDAC1 (top panel) and S215E phosphomimetic VDAC1 mutant (bottom panel). Recombinant wt-VDAC1 or S215E mutant was reconstituted into planar lipid bilayers. Representative current recordings with the corresponding amplitude histograms are shown. Current was monitored before and after addition of HKII in response to a −30 mV test potential during a 30-second duration. Downward deflections denote channel opening. The dashed blue lines denote 0-current levels.

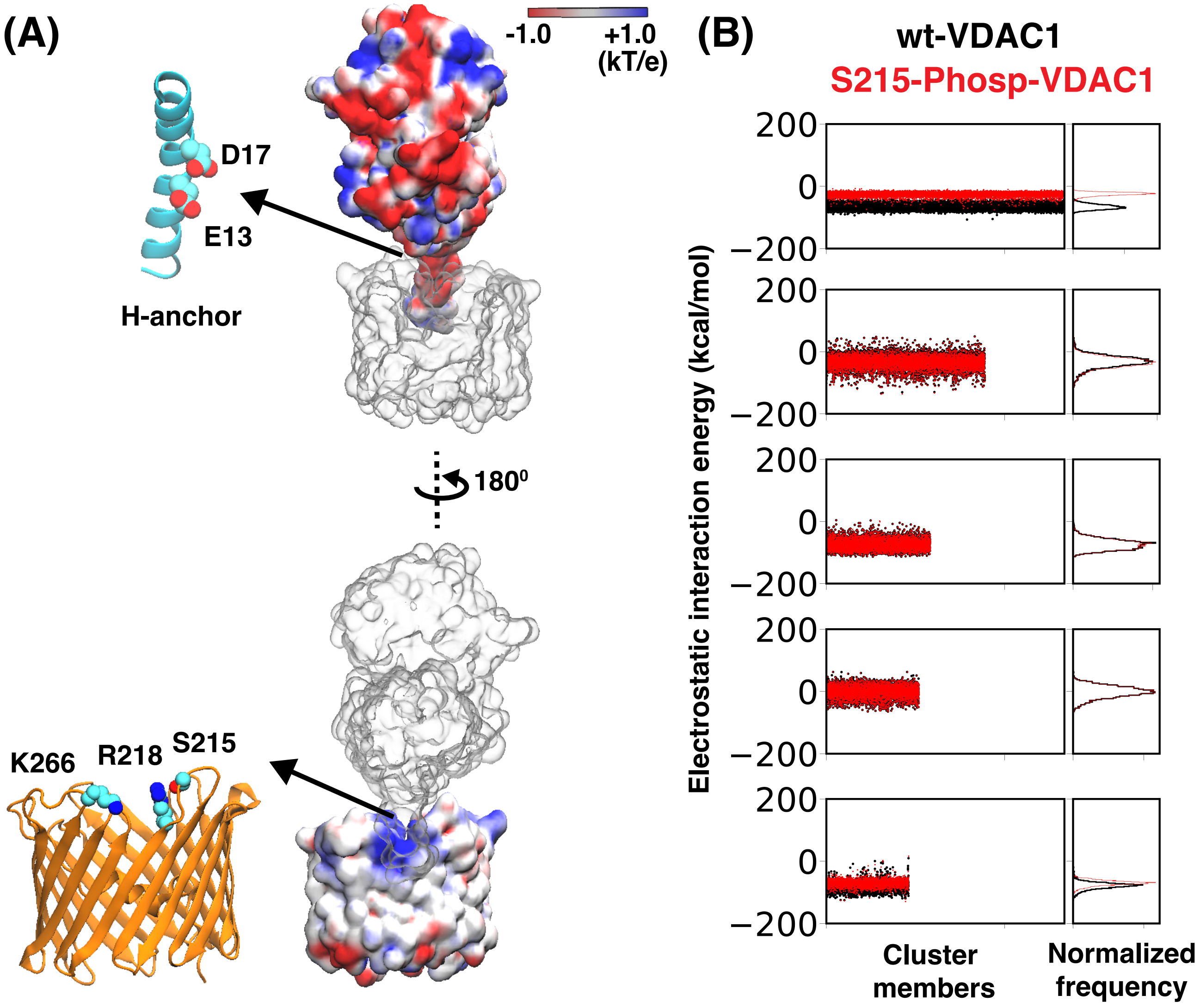


**Figure S6.** Proposed disruption mechanism of HKII/VDAC1 complex. (A) Electrostatic potential map of HKII-N and VDAC1 in HKV1 (generated using the Poisson-Boltzmann (PB) equation solver module in CHARMM-GUI[31–33]), highlighting the binding interface near H-anchor. The interface contains a electronegative surface in H-anchor and a electropositive surface in VDAC1. (B) Electrostatic interaction energy calculated between HKII and VDAC1 in two different states of VDAC1: wt-VDAC1 and S215-Phosp-VDAC1. The calculations were performed for all members of the five clusters (cluster 1 to 5) derived from BD simulation of wt-VDAC1.
